# Associations of prenatal and postnatal growth with insulin-like growth factor-I levels in pre-adolescence

**DOI:** 10.1101/144493

**Authors:** R Arathimos, C Macdonald-Wallis, CJ Bull, JMP Holly, E Oken, MS Kramer, N Gusina, N Bogdanovich, K Vilchuck, R Patel, R M Martin, K Tilling

## Abstract

**Background:** Rapid pre - and postnatal growth have been associated with later life adverse health outcomes, which could implicate (as a mediator) circulating insulin-like-growth-factor I (IGF-I), an important regulator of growth. We investigated associations of prenatal (birth weight and length) and postnatal growth in infancy and childhood with circulating IGF-I measured at 11.5 years of age.

**Methods:** We analysed 11.5-year follow-up data from 17,046 Belarusian children who participated in the Promotion of Breastfeeding Intervention Trial (PROBIT) since birth.

**Results:** Complete data were available for 5422 boys and 4743 girls (60%). We stratified the analyses by sex, as there was evidence of interaction between growth and sex in their associations with IGF-I. Weight and length/height velocity during childhood were positively associated with IGF-I at 11.5 years; associations increased with age at growth assessment and were stronger for length/height gain than for weight gain. The change in internal run-normalized IGF-I z-score at 11.5 years was 0.038 (95% CI -0.004,0.080) per standard deviation (SD) increase in length gain at 0-3 months amongst girls and 0.025 (95% CI - 0.011,0.060) amongst boys, increasing to 0.336 (95% CI 0.281,0.391;) and 0.211 (95% CI 0.165,0.256) for girls and boys, respectively, for growth during 6.5-11.5 years.

**Conclusion:** Postnatal growth velocities in childhood are positively associated with levels of circulating IGF-I in pre-adolescents. Future studies should focus on assessing whether IGF-I is on the causal pathway between early growth and later health outcomes, such as cancer and diabetes.

## Introduction

Insulin-like growth factor I (IGF-I) is an important regulator of childhood growth and development and a part of the insulin-like growth factor (IGF) system, also known as the IGF axis. The system is comprised of two ligands (IGF-I and IGF-II) and six high-affinity binding proteins (IGFBP-1-6). Growth throughout childhood is controlled by growth hormone (GH), the effects of which are mediated by IGF-I. In animal models, IGF-I deficiency caused by mutations in the *IGF-I* gene leads to both intrauterine and postnatal growth failure^(1)^, and in human studies, circulating IGF-I levels at birth are positively correlated with gestational age and birth weight standardized for gestational age^(2)^. Rate of weight gain between birth and 2 years of age is associated with IGF-I at 5 years of age^(3)^. Infants with higher circulating IGF-I have higher previous and subsequent growth rates, and IGF-I levels correlate with subsequent weight gain^(4)^. Weight and length/height at 1 year and 5 years of age have been associated with IGF-I levels at 25 years of age, indicating that IGF-I in adulthood may be related to growth in infancy^(5)^. Under-nutrition in utero may lead to reprogramming of the IGF-I axis^(6, 7)^. This hypothesis is supported by evidence from both experimental models^(8)^ and clinical paediatrics^(9)^.

In later life, higher circulating levels of IGF-I are associated with breast, prostate and colorectal cancers^(10–14)^, type 2 diabetes^(15, 16)^ and (albeit less consistently) coronary heart disease^(17–19)^. Anthropometric measures show robust associations with disease risk, for cancer, the risk of several cancers increases with height^(20)^ and leg length^(21)^, and IGF-I may act as an intermediate between growth and disease risk. Understanding the relationship between growth trajectories and IGF-I might inform understanding of disease mechanisms and assist in developing preventive or therapeutic interventions.

In the current study, we examine associations of anthropometric measures (weight and length/height) at birth and during childhood in relation to IGF-I levels measured in early adolescence. We use follow-up data from the Promotion of Breastfeeding Intervention Trial (PROBIT)^(22)^ cohort, a cluster-randomized trial investigating the long-term effects of a breastfeeding promotion intervention on later health and development.

## Methods

The design of PROBIT has previously been reported in detail ^(23)^. In brief, PROBIT enrolled 17,046 infants born in 1996-7 at 31 maternity hospitals in the Republic of Belarus. Trial inclusion criteria required infants to be healthy singletons born at ≥37 weeks, with birth weight ≥2500g and Apgar score ≥5 at 5 minutes, whose mothers initiated breastfeeding and had no illness that would interfere with breastfeeding. Over 95% of mothers in the maternity hospitals chose to initiate breastfeeding, and approximately 98% of those consented to participate in the trial.

The infants were followed up at 31 polyclinics (clinics providing outpatient paediatric healthcare) affiliated with the trial maternity hospitals at 1, 2, 3, 6, 9 and 12 months of age (n = 16,492; 96.7% response rate)^(22)^, at mean age 6.5 years (n = 13,889; 81.5% response rate)^(24)^ and at mean age 11.5 years (n = 13,879; 81.4% response rate)^(25)^.

### Ethics

PROBIT follow-up was approved by the Belarusian Ministry of Health and received ethical approval from the McGill University Health Centre Research Ethics Board; the Human Subjects Committee at Harvard Pilgrim Health Care; and the Avon Longitudinal Study of Parents and Children (ALSPAC) Law and Ethics Committee. A parent or legal guardian provided written informed consent in Russian at enrolment and at the follow-up visits, and all children provided written assent at the 11.5-year visit.

### Measurements of growth

Study paediatricians recorded clinically-measured weight and length during polyclinic visits at 1, 2, 3, 6, 9 and 12 months. At 6.5 years of age, study paediatricians measured standing height up to four times with a wall-mounted stadiometer (Medtechnika, Pinsk, Belarus), and weight with an electronic digital scale (Bella 840; Seca Corporation, Hamburg, Germany). At 11.5 years of age, study paediatricians measured standing height and weight using the same techniques and instruments as at the 6.5-year visit. At the 6.5-year study visit, paediatricians also abstracted from the polyclinic record all clinical measures of weight and height from 12 months of age to 6.5 years. Training, quality assurance and audit procedures have been described in detail previously^(26)^.

### Measurement of IGF-I

Whole blood samples were collected at the 11.5-year research visit by finger prick onto Whatman 903 filter-paper cards by the polyclinic paediatricians, after at least 8 hours of fasting^(27)^. The cards were then allowed to dry and subsequently stored in freezers at each of the 31 polyclinic sites at -20°C until transport to the laboratory at the National Mother and Child Centre in Minsk, where they were stored at -80°C. The samples for IGF-I were stored at -20°C for a median of 1.7 months (interquartile range [IQR], 1.0-5.1) and at -80°C for a median of 18.4 months (IQR, 13.3-21.6). Circulating IGF-I was quantified from a single 3mm diameter disc (approx. 3 μL of blood) per child after a single thaw, using the validated method of Diamandi et al^(28)^. Mean intra-assay coefficients of variation were 6%, 7%, and 9% for low, medium, and high IGF-I values, respectively; the respective inter-assay values were 8%, 12%, and 16%. The stability and recoverability of IGF-I from such dried bloodspot methods in PROBIT has previously been reported^(25)^.

IGF-I was assayed from 2 lots of reagents between January 2010 and November 2011 and, as we have noted previously^(25)^, assay kits of different lot numbers vary in measured IGF-I. To remove the potential effect of between-lot or between-run variations, we standardized values of IGF-I within each assay run (n=43) by computing z-scores ([IGF-I value - mean for each run]/ SD for each run).

### Other variables

Potential confounders considered in the analyses were maternal and paternal age at time of the birth of the index child, and maternal and paternal weight and height, as reported by the accompanying parent (usually the mother) at the 6.5 year follow-up. Mothers also reported maternal and paternal highest level of education and urban or rural residence at enrolment. Education was categorized into four levels as “complete university”, “advanced secondary/partial university”, “common secondary” and “incomplete secondary”. Confounders and other characteristics were tested for differences between excluded and included children using unpaired t-tests for continuous variables and chi-squared tests for categorical variables.

Tanner stage, a scale of physical maturation in children that defines development based on external primary and secondary sex characteristics, such as the development of the breasts in girls, genital (testicular volume) size in boys and development of pubic hair in both girls and boys^(29)^, were determined by direct physician examination at 11.5 years of age. Both Tanner stages for each sex (pubic hair and breast size for girls, pubic hair and genital size for boys) were used to adjust for pubertal onset in associations of growth rates with IGF-I stratified by sex.

### Statistical analysis

#### Effects of post-natal growth velocity

Multilevel models were fitted using MLwinN v2.26 to estimate each child's sex-standardised growth velocity from birth to 11.5 years. Linear splines were used to describe the shape of the growth trajectories, with knot points (indicating where the linear slope changes) based on a previously derived model of the dataset, positioned at 3 months, 12 months and 2.8 years^(30)^. The previous trajectories derived in PROBIT went up to only 60 months (5 years)^(30)^; the model was extended in the current analysis to include data at 11.5 years and thus included an additional knot point at 6.5 years. The average trajectories were allowed to differ by sex by including a fixed effect of sex and interactions by sex with each of the splines. Individual-level residuals were used to derive a predicted length and weight at birth, and rate of change in height and weight in each period of 0-3 months, 3-12 months, 1.1-2.8 years, 2.8-6.5 years and 6.5-11.5 years. These predicted growth rates were then used as the exposures in regression models with IGF-I at 11.5 years as the outcome.

All postnatal growth models used sex-standardised change in length/height or weight as the exposure. Robust standard errors were used to allow for polyclinic clustering. Model 1 adjusted for previous change(s) in length/height and exact age of the child at follow-up; model 2 adjusted additionally for parents' anthropometry, age and education level; and model 3 additionally for geographical location of the polyclinic (urban or rural; East or West of Belarus) and intervention arm. Stepwise adjustments for confounders in the 3 models ensured that parental covariates were adjusted for separately from confounding that relates to cohort design, such as trial arm and polyclinic location. As a sensitivity analysis for the growth models, a cross-sectional analysis at 11.5 years related observed height and weight at 11.5 years to measured IGF-I levels using linear regression. Additionally, a further sensitivity analysis was carried out using multiple imputation^(31, 32)^ to fill in missing values in covariates and IGF-I measurements. Imputation was performed in STATA 14.1 using the ICE process^(33, 34)^. We imputed 20 datasets to ensure estimate accuracy. Data was included for a total of 16,877 individuals that had attended at least 2 clinic visits, passed our quality-control (QC) and had recorded gender.

To determine whether the association between early growth and IGF-I was mediated by pubertal timing, growth rate models were adjusted for Tanner-based pubertal stage at 11.5 years of age in a separate model (Model 4). Although puberty cannot be a confounder of the growth-IGF-I association as it follows the previous measurement of growth, it may be a mediator of the growth-IGF-I association, particularly at 11.5 years, when many study children (especially girls) have entered puberty. To confirm that puberty was associated with in IGF-I in PROBIT, we also examined associations of Tanner stage with IGF-I at 11.5 years in girls and boys.

In models adjusting for interactions of weight or height with size-for-gestational-age categories, p-values were calculated by taking the difference in log-likelihood between the models adjusting vs not adjusting for the multiple interaction terms and tested using the chi-square statistic with 2 degrees of freedom.

#### Effects of size at birth

Categories of birth weight for gestational age were derived using a Canadian population-based reference^(35)^. Small-for-gestational-age (SGA) birth was defined as a birth weight <10th percentile for gestational age and sex, and large-for-gestational-age (LGA) birth was defined as a birth weight >90^th^ percentile for gestational age and sex. Infants with birth weights between these extreme percentiles were classed as appropriate for gestational age (AGA).

Linear regression models were fitted with birth weight-for-gestational-age categories as the exposure and IGF-I levels at 11.5 years as the outcome, accounting for clustering within the 31 polyclinics using robust standard errors. The reference category was AGA. Associations were estimated for both the whole sample and by sex, as growth rate residuals interacted with sex in their association with IGF-I.

## Results

Of the 17,046 infants originally recruited into PROBIT, a total of 10,165 (60%) with complete data on all exposures and confounders and with IGF-I measurements at 11.5 years were included in the analysis (Figure S1). All analyses were repeated on the full sample of 12,356 individuals with available IGF-I measures but with missing confounders. Little difference was found between the results of complete case analysis and the full dataset. Table 1 shows the baseline and follow-up characteristics of individuals in the analysis stratified by sex. Table S1 shows small differences between included versus excluded children in relation to geographical location and the educational attainment of the mothers and fathers.

**Table 1.** Baseline and follow-up characteristics of PROBIT children included in the current study by sex.

|  | Girls (N=4,743) | Boys (N=5,422) |
| --- | --- | --- |
| Height/length and weight - Mean (sd) |  |  |
| Birth length cm | 51.7 (2.1) | 52.4 (2.2) |
| Length/height at 3 months cm | 60.4 (2.3) | 61.6 (2.4) |
| Height at 11.5 years cm | 150.2 (7.9) | 148.7 (7.4) |
| Birth weight kg | 3.4 (0.4) | 3.5 (0.4) |
| Weight at 3 months kg | 5.9 (0.6) | 6.3 (0.7) |
| Weight at 11.5 years kg | 40.9 (9.5) | 40.6 (9.4) |
| Region* - N (%) |  |  |
| -East Urban | 1,462 (30.8) | 1,655 (30.5) |
| -East Rural | 755 (15.9) | 791 (14.6) |
| -West Urban | 1,147 (24.2) | 1,330 (24.5) |
| -West Rural | 1,379 (29.1) | 1,646 (30.4) |
| Maternal Education - N (%) |  |  |
| -Complete University | 686 (14.5) | 740 (13.7) |
| -Advanced Secondary/partial university | 2,452 (51.7) | 2,842 (52.4) |
| -Common or incomplete Secondary | 1,447 (30.5) | 1,693 (31.2) |
| -Incomplete Secondary | 158 (3.3) | 147 (2.7) |
| Fathers Education - N (%) |  |  |
| -Complete University | 614 (13.0) | 733 (13.5) |
| -Advanced Secondary/partial University | 2,288 (48.2) | 2,577 (47.5) |
| -Common or incomplete Secondary | 1,749 (36.9) | 2,005 (37.0) |
| -Incomplete secondary | 92 (1.9) | 107 (2.0) |
| Parental anthropometry - Mean (sd) |  |  |
| Mothers weight kg | 66.4 (12.6) | 66.4 (12.5) |
| Mothers height cm | 164.5 (5.6) | 164.9 (5.6) |
| Fathers weight kg | 79.6 (11.5) | 78.0 (11.5) |
| Fathers height cm | 176.0 (6.6) | 176.1 (6.6) |
| Mothers age | 25.2 (4.9) | 25.1 (4.9) |
| Fathers age | 27.5 (5.1) | 27.5 (5.1) |
| Tanner stage; pubic hair development - N (%) |  |  |
| Stage 1 | 1,448 (30.5) | 2,883 (53.2) |
| Stage 2 | 2,202 (46.4) | 2,152 (39.7) |
| Stage 3 | 971 (20.5) | 355 (6.6) |
| Stage 4 | 113 (2.4) | 27 (0.5) |
| Stage 5 | 9 (0.2) | 5 (0.1) |
| Tanner stage; female breast size/male genital size - N (%) |  |  |
| Stage 1 | 601 (12.7) | 727 (13.4) |
| Stage 2 | 2,453 (51.7) | 2,960 (54.6) |
| Stage 3 | 1,466 (30.9) | 1,546 (28.5) |
| Stage 4 | 208 (4.3) | 175 (3.2) |
| Stage 5 | 20 (0.4) | 14 (0.3) |
| IGF-I at 11.5 years ng/ml - Mean (sd) | 314.5 (124.9) | 244.8 (92.2) |

\* Regions of the Republic of Belarus divided in to East and West and by urban or rural housing tenure

No statistical evidence of interaction between sex and birth weight for gestational age on IGF-I was found (p=0.36) in fully-adjusted models. However, interaction tests of weight and height growth rate residuals with sex on IGF-I showed evidence of interaction at 2.8-6.5 years and 6.5-11.5 years for IGF-I (p<0.001 for both). Hence, all results are presented separately by sex.

The associations of sex-standardised length/height gain rate residuals with IGF-I at 11.5 years by sex are shown in Table 2. Neither birth length nor length gain velocity in the first 3 months was associated with IGF-I levels at 11.5 years. Associations of length/height gain with IGF-I were seen from 3 months onwards, however, and tended to be stronger in girls than in boys. The change in IGF-I z-score per SD change in length/height velocity between 2.8-6.5 years for girls was 0.236 (95% CI 0.196,0.276), and per SD change in length/height velocity between 6.5-11.5 years was 0.336 (95% CI 0.281,0.391). These respective coefficients in boys were 0.186 (95% CI 0.153,0.219) and 0.211 (95% CI 0.165,0.256). Associations between sex-standardised weight gain and IGF-I at 11.5 years, adjusted for previous measurements of height, are shown in Table 3. No associations were seen between birth weight and IGF-I levels at 11.5 years in boys; the change in IGF-I z-score per SD increase in birth weight was -0.007 (95% CI -0.044,0.032). An association was observed in girls, however: -0.065 (95% CI -0.115, -0.015). Amongst girls, no association was observed between weight gain velocity at 3-12 months of age and IGF-I (coefficient: 0.027; 95% CI: - 0.018,0.070), whereas a weak association was observed amongst boys (coefficient: 0.029; 95% CI 0.006,0.053). Weight gain rates between 2.8-6.5 years were not associated with IGF-I levels at age 11.5 years in boys; the changes in IGF-I z-score per SD increase in weight was 0.023 (95% CI -0.014,0.058). At 6.5-11.5 years there was only a weak association with IGF-I (coefficient: 0.036; 95% CI 0.003,0.070). The associations at the same two time-points were stronger in girls however: 0.101 (95% CI 0.045,0.158) and 0.140 (95% CI 0.088,0.193), respectively. A sensitivity analysis using multiple imputation to fill in missing data in 16,877 individuals revealed little difference in estimates when compared to the complete-case analysis (Table S7 and Table S8). Effect sizes for associations of length/height and weight growth rates with IGF-I in males increased marginally, whereas a small increase in effect sizes of weight growth rates in females was also evident. All effects were in the same direction as the complete cases analysis, except for birthweight and birth length/height in males which was reversed, with little evidence of an association in the imputed models for either.

**Table 2.** Associations of length/height gain rates (sex-standardized) with IGF-I z-scores at 11.5 years.

|  |  | **Model 1** | **Model 2** | **Model 3** |
| --- | --- | --- | --- | --- |
|  |  | Change in IGF-I z-score<br>per SD increase in height<br>(95% CI) | Change in IGF-I z-score<br>per SD increase in height<br>(95% CI) | Change in IGF-I z-score<br>per SD increase in height<br>(95% CI) |
| Girls | Birth | 0.014 (-0.016,0.044) | 0.004 (-0.028,0.037) | 0.007 (-0.021, 0.035) |
|  | 0-3 months | 0.05 (0.001,0.099) | 0.041 (-0.009,0.090) | 0.038 (-0.004,0.080) |
|  | 3-12 months | 0.132 (0.095,0.170) | 0.131 (0.093,0.170) | 0.13 (0.092,0.168) |
|  | 1.1-2.8 years | 0.038 (0.004,0.071) | 0.036 (0.003,0.069) | 0.032 (-0.003,0.066) |
|  | 2.8-6.5 years | 0.219 (0.183,0.256) | 0.236 (0.194,0.278) | 0.236 (0.196,0.276) |
|  | 6.5-11.5 years | 0.333 (0.279,0.387) | 0.337 (0.282,0.392) | 0.336 (0.281,0.391) |
| Boys | Birth | 0.002 (-0.031,0.034) | -0.005 (-0.038,0.027) | -0.005 (-0.037,0.026) |
|  | 0-3 months | 0.034 (-0.007,0.074) | 0.023 (-0.017,0.064) | 0.025 (-0.011,0.060) |
|  | 3-12 months | 0.119 (0.093,0.146) | 0.115 (0.086,0.144) | 0.113 (0.084,0.142) |
|  | 1.1-2.8 years | 0.032 (0.005,0.058) | 0.03 (0.004,0.056) | 0.032 (0.006,0.057) |
|  | 2.8-6.5 years | 0.176 (0.146,0.206) | 0.184 (0.153,0.216) | 0.186 (0.153,0.219) |
|  | 6.5-11.5 years | 0.208 (0.162,0.255) | 0.212 (0.165,0.259) | 0.211 (0.165,0.256) |

Three models with cumulative adjustments. Model 1 adjusted for age, birth length and previous measurement(s) of weight/height. Model 2 adjusted additionally for parents' anthropometry, age, and education. Model 3 additionally for region and treatment arm.

**Table 3.** Associations of weight gain rates (sex-standardized) with IGF-I z-scores at 11.5 years.

|  |  | **Model 1** | **Model 2** | **Model 3** |
| --- | --- | --- | --- | --- |
|  |  | Change in IGF-I<br>z-score per SD increase in<br>weight (95% CI) | Change in IGF-I<br>z-score per SD increase in<br>weight (95% CI) | Change in IGF-I<br>z-score per SD increase in<br>weight (95% CI) |
| Girls | Birth | -0.049(-0.107,0.01) | -0.062(-0.118,-0.007) | -0.065(-0.115,-0.015) |
|  | 0-3 months | 0.04(0.009,0.072) | 0.037(0.006,0.067) | 0.031(-0.002,0.063) |
|  | 3-12 months | 0.026(-0.015,0.067) | 0.022(-0.022,0.065) | 0.027(-0.018,0.07) |
|  | 1.1-2.8 years | 0.083(0.036,0.13) | 0.079(0.032,0.127) | 0.078(0.032,0.123) |
|  | 2.8-6.5 years | 0.11(0.051,0.17) | 0.106(0.047,0.164) | 0.101(0.045,0.158) |
|  | 6.5-11.5 years | 0.14(0.088,0.193) | 0.14(0.088,0.192) | 0.14(0.088,0.193) |
| Boys | Birth | 0.003(-0.038,0.043) | -0.008(-0.049,0.033) | -0.007(-0.044,0.032) |
|  | 0-3 months | 0.049(0.025,0.073) | 0.043(0.019,0.067) | 0.045(0.02,0.07) |
|  | 3-12 months | 0.037(0.013,0.061) | 0.031(0.006,0.057) | 0.029(0.006,0.053) |
|  | 1.1-2.8 years | 0.048(0.024,0.072) | 0.043(0.019,0.068) | 0.043(0.018,0.067) |
|  | 2.8-6.5 years | 0.036(0.002,0.07) | 0.022(-0.014,0.058) | 0.023(-0.014,0.058) |
|  | 6.5-11.5 years | 0.045(0.011,0.08) | 0.036(0.001,0.071) | 0.036(0.003,0.07) |

Three models with cumulative adjustments. Model 1 adjusted for age, birth length and previous measurements of weight/height. Model 2 adjusted additionally for parents' anthropometry, age, and education. Model 3 additionally for region and treatment arm.

Adjusting for Tanner stage at outcome attenuated some of the associations between height growth rate and IGF-I in girls (Table S6). Attenuation of associations of height growth rate between 2.8-6.5 years and 6.5-11.5 years with IGF-I for boys was also observed. For weight gain rates, adjustment for puberty affected the association with IGF-I only marginally between 2.8-6.5 years and 6.5-11.5 years in girls. However, adjustment for puberty increased the strength of the association somewhat between weight gain rates and IGF-I in boys.

Cross-sectionally, height and weight at 11.5 years were positively associated with IGF-I concentrations in both girls and boys (Table S2). Adjustment for puberty attenuated some of the effect sizes (Model 4 Table S2).

As expected, we observed a cross-sectional association between Tanner stage and IGF-I (Figure S2 & Table S5) with a 0.410 (95% CI 0.372,0.448) increase in z-score IGF-I per increase in Tanner stage for pubic hair development in girls and a 0.230 (95% CI 0.195,0.266) increase in z-score IGF-I per increase in Tanner stage for pubic hair development in boys. Adjusting for current height and weight at 11.5 years attenuated the effect sizes somewhat (Model 3 Table S5) but the effects persisted for both girls and boys.

No evidence for an association was found between birth weight-for-gestational-age categories and IGF-I z-score at 11.5 years in boys or girls. Compared to AGA births, the difference in IGF-I z-score in the fully-adjusted models (adjusted for age, birth-length, parent's anthropometry /age /education, region and treatment arm) for girls born SGA was 0.086 (95% CI -0.025, 0.197); for girls born LGA, the IGF-I z-score change was 0.004 (95% CI -0.096, 0.103). For boys born SGA, the difference in IGF-I z-score was -0.007 (95% CI -0.085, 0.071), and for those born LGA it was -0.034 (95% CI -0.127, 0.060) vs boys born AGA. Associations of birth weight-for-gestational-age categories with IGF-I levels at 11.5 years, and of interactions of birth weight-for-gestational-age categories with both weight and height, are shown in Table S3. No interactions were observed with weight or height in girls or boys.

## DISCUSSION

In this prospective analysis of over 10,000 Belarusian children, we observed positive associations of postnatal length/height and weight velocity with IGF-I levels at 11.5 years. These associations were stronger for growth in later than in earlier childhood, with little or no association of weight or length at birth with later IGF-I. In boys, weight gain during 2.8-6.5 years and 6.5-11.5 years was not strongly associated with later IGF-I. However, girls showed stronger associations of IGF-I with growth between 2.8-6.5 and 6.5-11.5 years of age. In general, associations began later in childhood in girls than in boys.

Observed associations attenuated once adjusted for Tanner stage. However, it is likely that puberty is a mediator of the relationship between growth and IGF-I, as rapid early growth may cause early onset of puberty, and IGF-I levels are higher during puberty. In this case, confounders of the puberty-IGF-I association will also become confounders of the growth-IGF-I associations, owing to collider bias^(36, 37)^ induced by conditioning on pubertal stage. Furthermore, recent evidence from animal models suggests IGF-I is involved in the regulation of the onset of puberty^(38)^. The biological pathways involved are likely complex, with many feedback loops involving both IGF-I and puberty onset. Further work is required on the subject before conclusions can be drawn about the exact role of IGF-I. A distinction should also be made between a statistical mediator and biological mediator. Whereas a confounding “third variable” could help explain an association between two other variables (which could be said to ‘cause' the association), it may have nothing to do with the underlying biological cause or biological pathway. Causality and mediation can have different inferences when considered statistically vs biologically.

Additionally, substantial variability has been observed between Tanner stages and other methods of assessing maturation or growth, such as peak height velocity (PHV), suggesting that Tanner stages may be an imprecise measure of pubertal onset^(39)^. This may lead to incomplete adjustment for puberty. The degree to which onset of puberty explains the observed associations may be studied through mediation analysis^(40)^ in future work.

### Comparisons with previous studies

IGF-I has an important role in regulating childhood growth. In agreement with our findings in PROBIT, children from the Avon Longitudinal Study of Parents and Children (ALSPAC) cohort who had greater gains in weight during early childhood had higher IGF-I levels at 5 years^(3)^. Specifically, an association between weight gain rate between 0–2 years and IGF-I at 5 years was observed in ALSPAC. We observed similar associations in PROBIT between weight gain rate and IGF-I at 11.5 years for boys at 0-3 months, 3-12 months and 1.1-2.8 years. However, for girls the same associations were not statistically significant at 0-3 months or 3-12 months. Instead, girls showed a strong association of IGF-I with weight gain rate in later childhood, i.e., at ages 2.8-6.5 years and 6.5-11.5 years. Our findings suggest substantial differences in magnitude of association between growth and IGF-I in girls vs boys. In ALSPAC, sex also appeared to modify the associations of IGF-I with leg growth, but the association was stronger in boys than in girls (p for interaction = 0.041)^(41)^. In relation to programming effects in ALSPAC, IGF-I levels at 5 years were inversely related to birth weight^(3)^. In PROBIT, a similar inverse association between birth weight and IGF-I at 11.5 was found only in girls. No associations with size-for-gestational-age categories were found, however, suggesting that the association with crude birth weight in ALSPAC and PROBIT may have been driven by gestational age at birth. A study of 951 individuals followed from birth reported an association of IGF-I in early adult life (mean age of 25 years) with height at 25 years of age and patterns of childhood growth but not with anthropometric measures at birth (including birthweight and birth length)^(5)^. Similarly, we observed that birth length was not associated with IGF-I levels at 11.5 years in boys or girls and that birth weight was associated with IGF-I in girls (but not boys).

A historical cohort study based on 65 years of follow-up of 429 individuals found that neither height nor leg length in childhood or adulthood was associated with circulating levels of IGF-I, IGF-II, or IGFBP-2 measured in late adulthood^(42)^. However, in the latter study, associations between components of height and IGF-I were all positive with wide confidence intervals, and thus are compatible with findings in PROBIT. Differences in ages between the two samples, however, make it difficult to draw conclusions, as IGF-I patterns change throughout life and many effects may have resolved by adulthood.

### Strengths and limitations

Follow-up rates in PROBIT were high, minimising selection bias. However almost 28% of study children did not have measured IGF-I at 11.5 years. IGF-I was missing in individuals that did not attend clinic follow-up at 11.5 years. IGF-I was also not measured in individuals who did not provide enough blood sample for the assay or were excluded for technical reasons during quality control. The level of missing data on IGF-I could lead to biased associations if the likelihood of missing IGF-I is related to underlying IGF-I, conditional on the variables in the model, which we consider unlikely. A sensitivity analysis for the growth models using multiple imputation did not reveal substantial differences in estimates when compared to the complete cases analysis, indicating that any such bias is unlikely. The study sample population was sufficiently large to detect modest associations and interactions. Multiple measures of growth were available across infancy and childhood, and appropriate statistical methods were used to relate growth to IGF-I.

Limitations include the fact that weight and length measurements in infancy were not taken according to a standardised protocol, which could lead to non-differential measurement error and thereby attenuate associations. Uncertainty in the modelled estimation of the growth trajectories was not accounted for, and thus the width of the confidence intervals may be underestimated. Measurement of IGF-I levels by dried blood spot may have led to measurement error; however, any error is again likely to be non-differential with respect to IGF-I and would therefore bias associations towards the null. Finally, the setting in Belarus may limit generalizability to other settings.

## Conclusion

Postnatal growth velocity in Belarusian children was associated with higher circulating IGF-I at 11.5 years of age. The strength of the growth rate association with later-life IGF-I differed by sex. These results support the association between later-life IGF-I and childhood growth previously reported and support the hypothesis that IGF-I levels in childhood might mediate an association between anthropometric measures and disease in later-life. Future studies should assess whether IGF-I mediates the effect of early growth on later health.

